# From cell types to cell states: adapting foundation models for cellular plasticity

**DOI:** 10.1101/2024.08.16.608311

**Authors:** Shi Pan, Eloise Withnell, Cenk Celik, Maria Secrier

## Abstract

Single-cell foundation models (scFMs) have been widely adopted for cell type annotation, yet their suitability for modelling cellular plasticity, where cells transition along continuous, context-dependent state trajectories, remains unclear. Here, we systematically benchmark state-of-the-art scFMs against conventional machine learning and bioinformatics approaches on epithelial-mesenchymal plasticity (EMP), a prototypical plastic cellular process. We show that naïve fine-tuning of scFMs often fails to resolve intermediate states and is strongly influenced by tissue- and stimulus-specific signals. We propose a parameter-efficient dual-task adaptation strategy for EMP foundation models (EMP-FM) that combines discrete classification with pseudotime-guided regression, which improves cell state resolution in controlled settings (up to 85% AUROC), but remains sensitive to domain shifts. Across diverse *in vitro* and *in vivo* datasets, scFMs do not consistently outperform conventional methods, which often achieve comparable performance with lower complexity. Together, our results delineate both the potential and current limitations of scFMs for modelling cellular plasticity and support their complementary use alongside established bioinformatics approaches.

## INTRODUCTION

Single cell omics technologies have made it possible to characterise cellular identity and behaviour at unprecedented resolution. While much of the field has focused on cell type classification, many biologically important phenomena manifest not as discrete identities but as continuous, context-dependent cell states. Cell <u>types</u> represent relatively stable, discrete cellular identities (e.g., T cells, B cells, fibroblasts) that can be effectively distinguished through categorical classification approaches. In contrast, cell *<u>states</u>* (e.g. stemness, metabolic rewiring, activation) reflect dynamic, context-dependent behaviour that cells adopt along a biological continuum. A classical example is the epithelial-to-mesenchymal transition (EMT), a developmental process in which epithelial cells gradually acquire mesenchymal characteristics, and which is frequently co-opted in cancer to promote invasion and metastasis^1,2^. EMT and other plastic processes, such as differentiation and immune activation, are characterised by a continuum of transformation rather than stable, well-separated transcriptional categories^3^, are stimulus-dependent and may lack clear boundaries. As a result, they pose fundamentally different computational challenges.

Traditional bioinformatics approaches, typically relying on canonical markers (e.g. E-cadherin or vimentin^4–6^ for EMT) or limited gene signatures^7,8^, and conventional machine learning approaches designed for discrete classification offer valuable insights but are inherently ill-suited for modelling the fluid, continuous nature of cellular plasticity. As cancer cell trajectories vary depending on the tissue and external stimuli^9^, these strategies struggle to resolve subtle intermediate states or to generalise across experimental conditions. Furthermore, continuous processes often involve long-range or nonlinear regulatory dependencies^10^ that are not easily captured by linear models or shallow classifiers, limiting our ability to develop unified frameworks to characterise cellular plasticity in diverse systems and beyond tightly controlled *in vitro*/*in vivo* settings.

Single cell foundation models (scFMs) like scBERT^11^, scGPT^12^, Cell2Sentence^13^, GenePT^14^ or scFoundation^15^, have introduced a powerful new paradigm that holds promise in addressing such challenges. These models are trained on extremely large and diverse scRNA-seq corpora, leveraging transformer architectures^16–18^ that enable them to encode broad transcriptional programmes, learn long-range gene-gene dependencies and, in principle, provide transferable representations for downstream tasks^19^. However, current scFMs have been predominantly developed and evaluated for discrete annotation tasks, especially cell type classification. This creates a fundamental mismatch when applied to continuous biological processes like EMP: the learned representations tend to prioritise strong sources of variation, such as tissue of origin or stimulus, over the more subtle, dynamic signals that define cell state transitions. Furthermore, recent evaluations also reveal significant limitations of these models in a zero-shot cell-type classification scenario, with scFMs often underperforming compared to simpler baselines^20^. However, the potential of fine-tuning scFMs for cell state classification remains insufficiently explored in systematic settings.

These limitations raise two important questions. First, can existing scFMs be effectively adapted to model continuous cellular processes, or do they require fundamentally different training objectives and architectures? Second, what design principles are needed for foundation models to capture dynamic transcriptional changes that unfold along biological continua? To explore these questions, we focus on epithelial–mesenchymal plasticity (EMP) as a representative system. EMP provides a well-characterised yet challenging example of a continuous transition involving heterogeneous intermediate states, strong context dependency, and substantial variability across cancer types and stimuli. Importantly, it offers an experimentally grounded testbed for assessing the strengths and weaknesses of current scFM paradigms when repurposed for cell state modelling.

By systematically benchmarking conventional approaches and several state-of-the-art scFMs on modelling EMP across multiple cancer types and under different stimuli, we uncover fundamental limitations in how pre-trained scFMs represent cell states and demonstrate why naïve fine-tuning is insufficient to overcome these issues. To address this, we develop a parameter-efficient dual-path adaptation framework that reshapes the pretrained feature space to better capture context-dependent trajectories, while preserving the underlying structure learned during large-scale pretraining. Our results highlight both the promise and the current limitations of using foundation models for continuous cell-state modelling, and outline conceptual and methodological directions for future scFM development.

## RESULTS

### EMP-FM, a foundation model that predicts EMT states in single cell cancer data

To facilitate a better understanding of cellular plasticity, we explored whether a pre-trained foundation model can quantify EMP from scRNA-seq data. We introduce EMP-FM, a fine-tuned scFM designed to probe whether targeted architectural and training choices using both a classification and a regression strategy can improve the representation of cell states along the EMT continuum. To define the ground truth transcriptional states that cells occupy as they undergo EMT in different cancer tissues, we employed data from Cook and Vaderhyden^9^. This study encompasses single cell RNA sequencing from breast (MCF7), lung (A549), prostate (DU145) and ovarian (OVCA420) cancer cell lines undergoing EMT, profiled at 0 days, 8 hours, 1 day, 3 days and 7 days after stimulation with TGFβ, TNF or EGF. We labelled cells sequenced at day 0 as epithelial (E), and ulterior states as hybrid E/M (EM1 at 8 hours, EM2 at 1 day, EM3 at 3 days) and mesenchymal (M, 7 days), respectively (**Supplementary Figure 1**). This experimental setup provides clean transcriptional profiles for distinct states along the EMT continuum.

The EMP transcriptional logic can be learned from the experimentally measured longitudinal profiles of this dataset. We followed the embedding strategy in other pre-trained scFMs^11,12^ and designed a two-stage dual path pipeline consisting of extra residual connections to enhance the raw gene expression signal, a Parameter-Efficient Fine-Tuning (PEFT)^21^ module and a Mixture of Expert (MoE)^22^ architecture with: (1) a classification head to learn discrete cell state labels, and (2) a regression head using the pseudotemporal ordering calculated by psupertime^9^ (**Figure 1a-b and Methods**). The dual modelling classification/regression strategy addresses the inherent challenge that EMP represents a continuous biological process that cannot be fully captured by discrete classification alone, thus transforming the pre-trained scFM from a general cell type classifier to a specialized cell state predictor capable of capturing fine-grained EMP cell state transitions.

**Figure 1:**
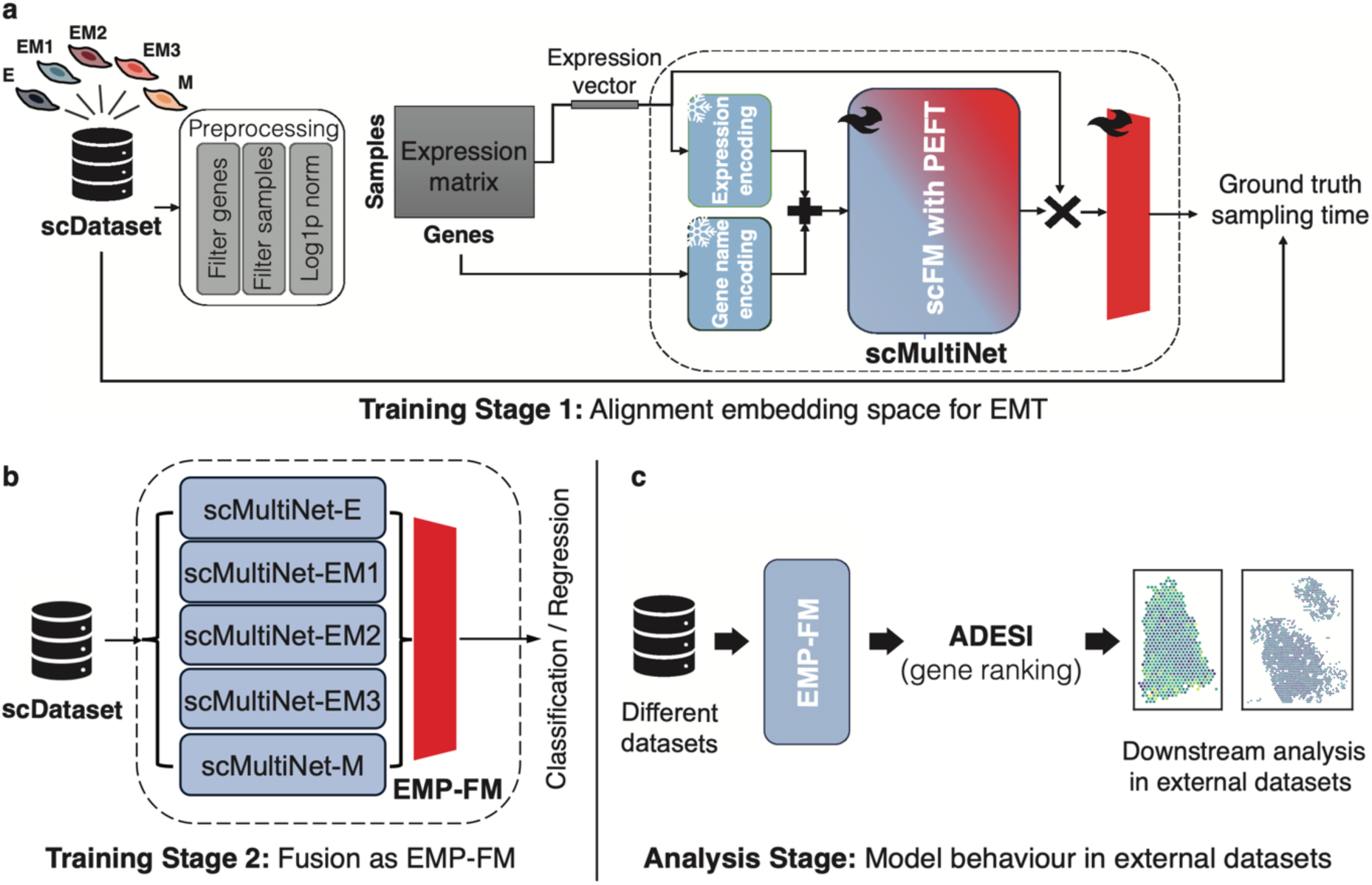
**EMP-FM pipeline in scRNA-seq data**. **(a)** The first training phase involves binary classification training for each EMT time point using the PEFT method. This allows the original pre-trained model’s feature space to better represent EMT. **(b)** The final model is obtained by fusing the five binary classification models for the individual states (E, EM1-3, M). **(c)** Downstream analysis employs ADESI to rank genes that contribute significantly to the model based on attention scores and gene expression.

The feature space from the pre-trained backbone of scBERT^11^ was modified to learn a generalisable EMP process across different tissues, and each expert network (named scMultiNet) focused on an individual target cell state across multiple tissues. Finally, we add an interpretability module to allow the exploration of the gene regulatory programmes underlying these states (**Figure 1c**).

### EMP-FM behind the scenes - gradual learning of EMP states

Understanding the patterns learned by EMP-FM is a crucial step in assessing whether pre-trained scFMs can prioritise subtle process-specific transitions like EMP over dominant signals such as tissue type. We explored the learned latent space evolution through five embedding stages using principal component analysis (PCA) (**Figure 2a)**: the original data distribution, the feature space of the data after being processed by the original scBERT, the feature space generated by scBERT after PEFT, the feature space of our proposed scMultiNet, and finally, the feature space resulting from the EMP-FM fusion. In the raw gene expression data, cells clustered by cancer cell line rather than experimental timepoints, demonstrating how tissue-specific signatures mask EMP patterns. Higher-order principal components (**Supplementary Figure 2a**) showed intermixing of EMP states rather than clear EMP trajectories, confirming that EMP states are inseparable in the raw data. While the pre-trained scBERT reduced cell line batch effects, it failed to separate EMP timepoints, with cells remaining extensively intermixed (**Figure 2a**). This exemplifies how pre-trained scFMs, despite zero-shot cell type classification capabilities, struggle to capture subtle variations in biological processes across tissues and stimuli. Our proposed scMultiNet achieved more distinct boundaries and tighter clustering between EMP categories, while the final EMP-FM model showed distinct separation of all EMP states, demonstrating successful capture of EMP-related features (**Figure 2a**). Remarkably, although the model was trained using discrete labels <u>without explicit temporal</u> <u>information</u>, it also successfully captures the sequential nature of the EMT process, with cells naturally arranging themselves from top right to bottom left (blue to orange) on a chronological EMT continuum in the latent space (**Figure 2a**, also shown with PHATE^23^ and Diffusion Pseudotime (DPT)^24^ analysis in **Supplementary Fig 2b**). This demonstrates the model’s ability to learn the underlying dynamics from the data.

**Figure 2:**
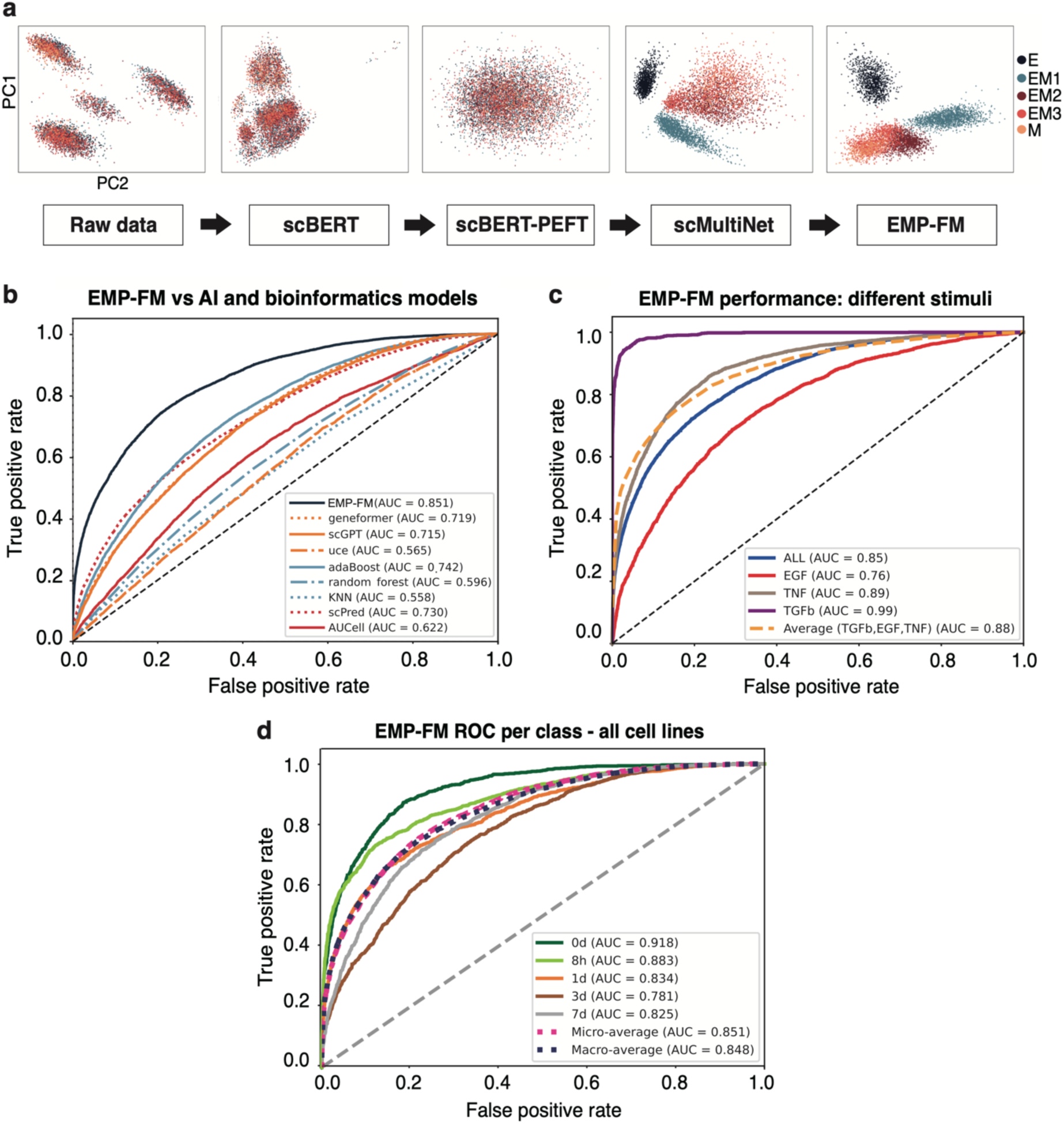
EMP-FM embedding space visualisation and benchmarking. **(a)** The changes in embedding space are shown through PCA projections from various models, highlighting the transformation of raw data into refined clusters following the application of scBERT, PEFT, scMultiNet and EMP-FM. Every point corresponds to a single cell that has been profiled. The five colours indicate the ground truth EMP states. **(b)** Comparison of EMP-FM with state-of-the-art foundation models including scGPT, Geneformer and UCE, conventional machine learning and bioinformatics methods. **(c)** EMP-FM performance under different EMT-inducing stimuli. ROC curves are shown for models of TGFβ-, TNF- and EGF-induced EMT, alongside an all-in-one model (ALL), with the average performance across the three stimulation types displayed as dotted curve. **(d)** EMP-FM-ALL performance across different timepoints and EMT states: epithelial (E), early EMT (EM1), intermediate EMT (EM2), late EMT (EM3), and mesenchymal (M).

### Benchmarking of EMP-FM

Having established that EMP-FM can learn meaningful EMT representations across conditions, we benchmarked its classification performance against state-of-the-art scFMs, conventional machine learning and bioinformatics methods. All methods were trained on the same 80% randomly sampled single cells across distinct EMT-inducing stimuli (TGFβ, TNF, EGF) and tested on the remaining 20%. EMP-FM achieved 85% AUROC, outperforming scFMs with standard fine-tuning: scGPT^12^ (72%), Geneformer^25^ (73%), and Universal Cell Embedding (UCE)^26^ (56.7%) (**Figure 2b**, **Supplementary Table 1**). EMP-FM’s performance also exceeded that of conventional machine learning approaches including K-Nearest Neighbours (KNN, 57%), random forest (75%) and AdaBoost (83%) (**Figure 2b, Supplementary Table 1**). Among specialised bioinformatics methods, scPred^27^, a supervised method to discretise cell types/states through feature selection on highly variable genes, showed similar performance (74%) to some of the other fine-tuned scFMs, while AUCell^28^, which assigns a continuous score per cell based on the expression of a given gene set, achieved a lower performance of 62%. scType^29^, a cluster annotation method based on known cell markers, could only distinguish epithelial and mesenchymal states, mis-annotating most hybrid states (**Figure 2b, Supplementary Figure 3**). Overall, EMP-FM achieved higher classification performance compared to both traditional methods and state-of-the-art scFMs in capturing targeted EMP cell states in this controlled setting.

We also built stimulus-specific EMP-FMs, which only adapted the classification head in the fusion network and were trained exclusively on individual stimulation conditions across all tissues (TGFβ, TNF, EGF). EMP-FM-TGFβ achieved 99% AUROC, while EMP-FM-TNF and EMP-FM-EGF reached 89% and 76% AUROCs, respectively (**Figure 2c, Supplementary Figure 4, Supplementary Tables 1-2**). Our results are in agreement with previous reports^9^, suggesting that the canonical EMT inducer TGFβ drives coordinated transcriptional changes delineating a comparatively more deterministic process^30–32^. Capturing EGF-induced transitions was more challenging, potentially due to increased transcriptional heterogeneity^33^, while in the case of TNF-induced EMT the models may struggle to capture complex pathway synergies required for stable EMT phenotypes^34^.

All EMP states showed good discrimination with AUROCs greater than 78% in the EMP-FM-ALL model and greater than 98% in the TGFβ model (**Figure 2d**). Both TNF and EGF models performed best at the 3-day timepoint (EM3: 98% and 85% AUROC, respectively), showing superior discrimination of intermediate and late EMT compared to early 8-hour transitions which are likely less stable (**Supplementary Table 2**). Notably, here each model captured stimulus-specific programmes, as applying them to the other stimulus scenarios decreased classification performance significantly (**Supplementary Table 3**).

These results show that generic scFMs, although minimally reliant on preprocessing or curated gene lists, do not consistently outperform conventional machine learning or bioinformatics approaches for cell state classification, with methods such as AdaBoost and scPred often achieving comparable or superior performance. Carefully designed, parameter-efficient adaptations have can substantially improve scFM performance, but remain sensitive to context and experimental conditions.

### Intra-dataset and domain-shift validation

To evaluate the generalisability and limitations of scFMs for cell state learning tasks, we conducted validation in four EMP-centric experimental settings: mesenchymal-to-epithelial reversal experiments from the same study by Cook and Vanderhyden, independent cell line datasets under different experimental designs, genetically engineered mouse model (GEMM) data, and human *in situ* datasets (spatial transcriptomics).

Cook and Vanderhyden^9^ tested two EMT reversal scenarios in their study: (1) where the EMT-inducing stimulus was removed from mesenchymal cells, and (2) where mesenchymal cells were treated with EMT inhibitor LY364947 (targeting TGFΒR1). Both scenarios should induce partial epithelial reversion, which allows us to question whether the model has learned EMP biology rather than memorising time-point associations. Neither scenarios would be expected to show a complete reversion to the 0-day state, or to follow a similar trajectory from M to E as from E to M. EMP-FM confirmed a significant reduction in EMT (Wilcoxon rank-sum test p < 0.001) following both inhibitor treatment (**Figure 3a-b**) and stimulation removal (**Figure 3c, Supplementary Figure 5a**). These results demonstrate that scFMs can learn meaningful biological transition patterns rather than noise or batch effects, with successful bidirectional prediction validating its understanding of continuous EMT regulatory logic across distinct experimental perturbations. However, not all EMP states remained consistently present during the reversal process (**Figure 3d, Supplementary Figure 5b**), likely reflecting subtle differences in dominant cell states and cancer trajectories under varying conditions, as posited by Cook and Vanderhyden^9^. This is a common challenge in cell state tasks that most classification-only scFM frameworks cannot adequately address. Our proposed dual-path architecture incorporating a regression element can overcome this by assigning continuous EMT scores, which revealed the expected downward trends in this experiment (**Supplementary Figure 5a**).

**Figure 3:**
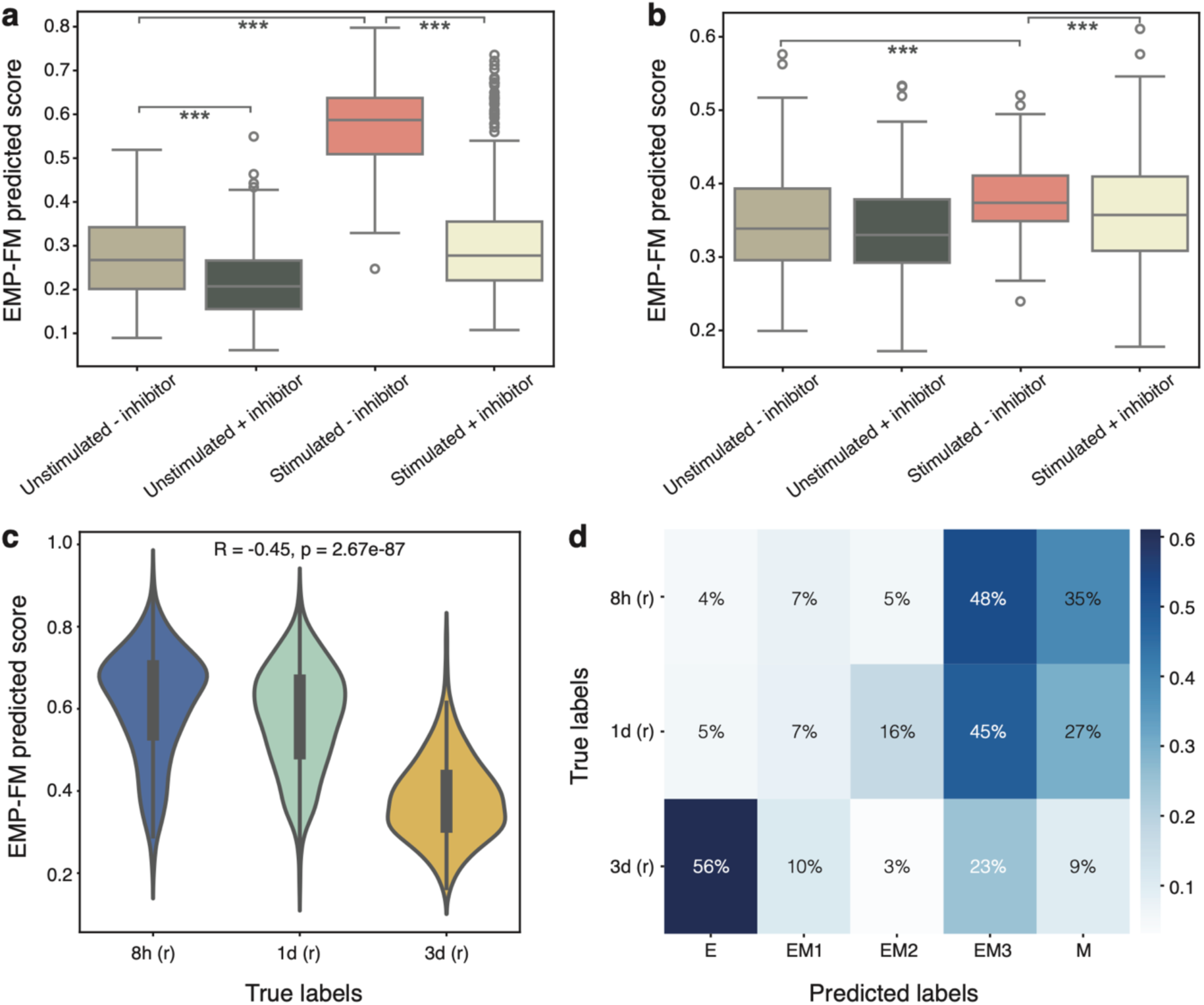
Model validation in EMT inhibitor and stimulus removal data in the original study. (a-b) Predicted EMP-FM-TGFβ scores in A548 (a) and MCF7 (b) cell lines that are either unstimulated or stimulated with TGFβ, and either treated (+inhibitor) and untreated (-inhibitor) with a TGFBR1 inhibitor. (*** p-value <0.001, ** <0.01, *<0.1). EMT scores are significantly lower after the addition of the inhibitor and in the unstimulated versus stimulated condition, as expected. **(c)** EMP-FM-TGFβ regression score distribution for the TGFβ stimulus removal data across all cancer cell lines, sequenced at 8h, 1 day and 3 days post-removal, respectively. The scores decrease with increasing time after removal, as expected. The Spearman correlation coefficient (R) and p-value are reported. **(d)** Confusion matrix of predicted cell states for the cells with TGFβ removal (averaged across all cancer cell lines) from the EMP-FM-TGFβ model.

We further explored the potential and limitations of scFMs in tracking EMT progression *in vitro* using external cell line scRNA-seq datasets: MCF10A breast epithelial cells treated with TGFβ from Paul et al^35^, and MCF10A cells undergoing spontaneous or TGFβ-induced EMT from McFaline-Figueroa et al^36^. Compared to our training data, the Paul et al^35^ dataset used the same TGFβ stimulus but with denser temporal sampling, while McFaline-Figueroa et al^36^ provided a different experimental design that captured both spontaneous and TGFβ-induced EMP processes at two time points, before and after EMT. As directly applying the EMP-FM classifier head to these datasets faced challenges due to mismatched class numbers between training and testing datasets, we generated continuous scores instead to provide more appropriate evaluation metrics and constructed a pseudo confusion matrix to demonstrate limitations of scFMs. In the Paul et al^35^ dataset, EMP-FM-TGFβ showed consistent EMT score increases across T0-T4 timepoints, as expected, with T4-T7 reaching steady state (**Figure 4a-b, Supplementary Tables 4-5**) and outperformed differentially expressed gene (DEG) signatures that failed to capture T3 transitions (**Supplementary Figures 6, 7a**). In the McFaline-Figueroa et al^36^ dataset, EMP-FM-TGFβ specifically captured TGFβ-driven but not spontaneous EMT increases, while EMP-FM-TNF, EMP-FM-EGF and all-in-one models detected increases in both conditions, suggesting spontaneous EMT resembles TNF/EGF-induced pathways (**Supplementary Figures 8, 9a, Supplementary Table 5**). However, this does not indicate that scFMs simply outperform conventional approaches; conversely, in some cases we observed that traditional bioinformatics methods outperform scFMs, particularly for unseen experimental designs (i.e. spontaneous EMT scenarios, **Supplementary Table 5**).

**Figure 4:**
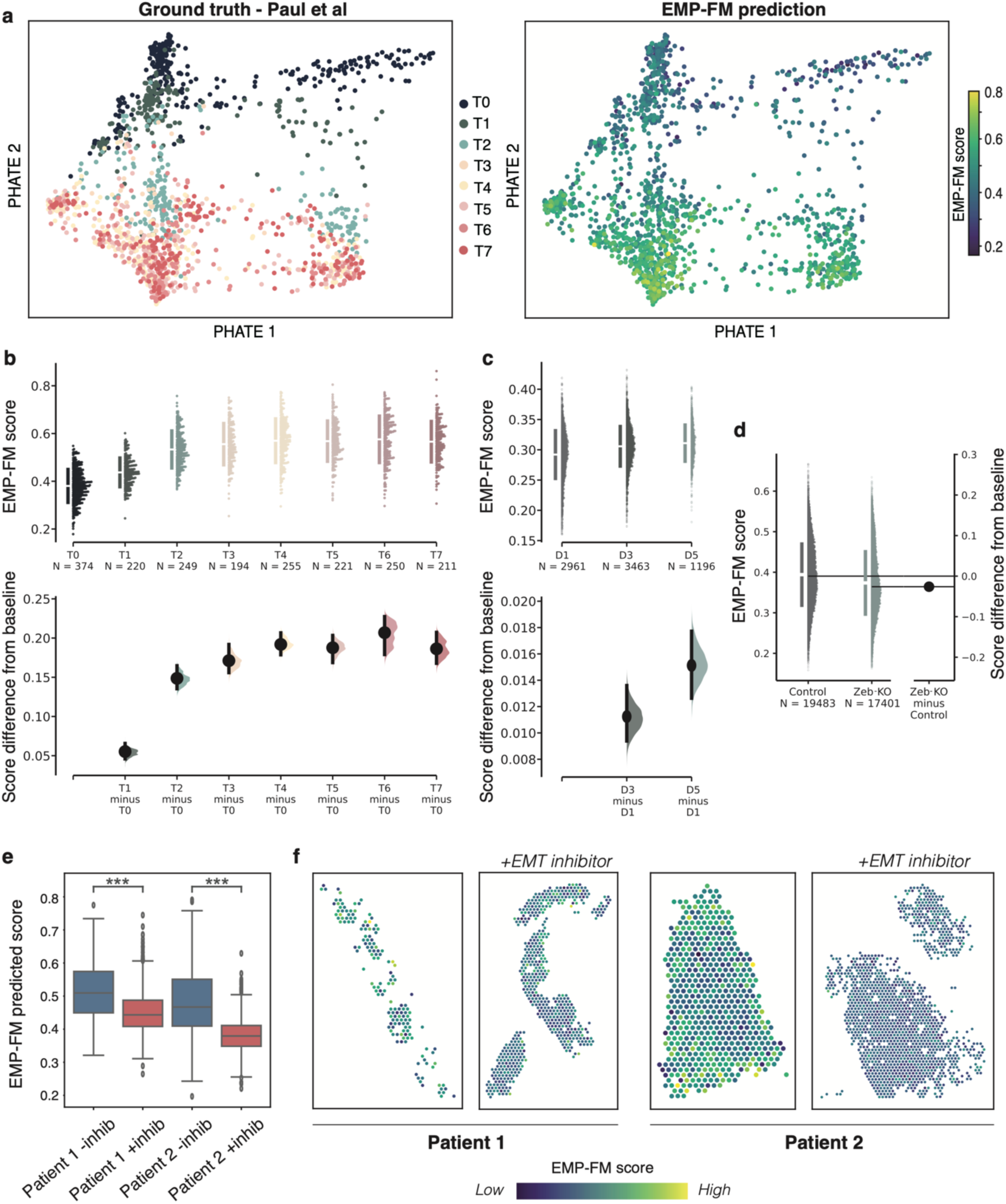
Validation of the EMP-FM model in external datasets. (a-b) Model validation in MCF10A cancer cell lines undergoing EMT (Paul et al^35^): **(a)** PHATE visualisation of individual cell projections coloured by ground truth sampling time (left panel, time points measured T1-T7) and scores predicted by EMP-FM (right panel, higher values indicate a more mesenchymal state); **(b)** Distribution of EMP-FM scores predicted for individual cells at the 7 time points (top) and differences in scores compared to the T0 baseline (bottom). Increases at all time points are significant (Kruskal-Wallis test p<<0.001, see statistics in Supplementary Table 4). **(c)** Model validation in C3(1)-Tag mouse breast cancer cells tracked during hybrid E/M transition (Grasset et al^37^): Distribution of EMP-FM scores predicted for individual cells at day 0 (12h), day 3 and day 5 (top) and differences in scores compared to the day 0 baseline (bottom). Increases at both time points are significant (Kruskal-Wallis test, see statistics in Supplementary Table 4). **(d)** Model validation in mouse basal mammary epithelial cells tracked upon Zeb1 knockout (Han et al^38^): Distribution of EMP-FM scores predicted for individual cells before and after Zeb1 knockout (left) and score difference compared to control (right). Increases after knockout is significant (Kruskal-Wallis test, see statistics in Supplementary Table 4). **(e-f)** Model validation in an external EMT inhibitor (Netrin-1) dataset from Cassier et al^39^: **(e)** Score distribution across endometrial spatial transcriptomic slides before (-inhib) and after (+inhib) treatment with the EMT inhibitor; **(f)** Visualisation of EMP-FM predicted regression scores across the spatially profiled slides before and after EMT inhibition in Patient 1 (left) and Patient 2 (right).

We next extended our validation to experiments performed in GEMMs to evaluate the generalisation capabilities of scFMs with domain shifts *in vivo* in two scenarios: (1) breast cancer cells tracked as they become increasingly invasive (thus activating EMT) at 12 hours, 3 days, and 5 days from Grasset et al^37^; and (2) knockout of a key EMT regulator, Zeb1, in basal mammary epithelial cells from Han et al^38^. In the cell invasion dataset, EMP-FM captured the expected gradual EMT progression at days 3 and 5 (**Figure 4c**), while differentially expressed gene signatures showed bimodal distribution at day 3 and unexpected decreases at day 5 (**Supplementary Figure 9b**). In the Zeb1 knockout setting, we observed decreased EMT scores consistent with expected EMT attenuation following Zeb1 loss (**Figure 4d**), whereas gene signatures incorrectly predicted EMT increases (**Supplementary Figure 9c**). However, despite incorporating broader gene associations, EMP-FM showed comparable performance to conventional machine learning and bioinformatics methods in several instances (**Supplementary Table 4-5**). Furthermore, training data composition significantly influenced the scFMs’ predictive performance across different GEMM datasets, while traditional methods provided relatively stable results with substantially lower computational overhead.

Finally, EMP-FM was further evaluated using spatial transcriptomics (ST) data from Cassier et al^39^, profiling endometrial tumours from two patients before and after Netrin-1 EMT inhibitor treatment. It is worth noting this is a challenging test setting due to differences between scRNA-seq and ST data and distinct cancer tissue. Without additional processing steps, our model predicted decreased EMP scores post-treatment in both cases (**Figure 5e**), with reductions visible throughout tissue slides (**Figure 5f**). The fact that our model is able to capture this in endometrial cancer, which is not a cancer type it was originally trained on, suggests that aspects of the learned EMP-related signal may transfer beyond the cancer types represented in the training data.

**Figure 5:**
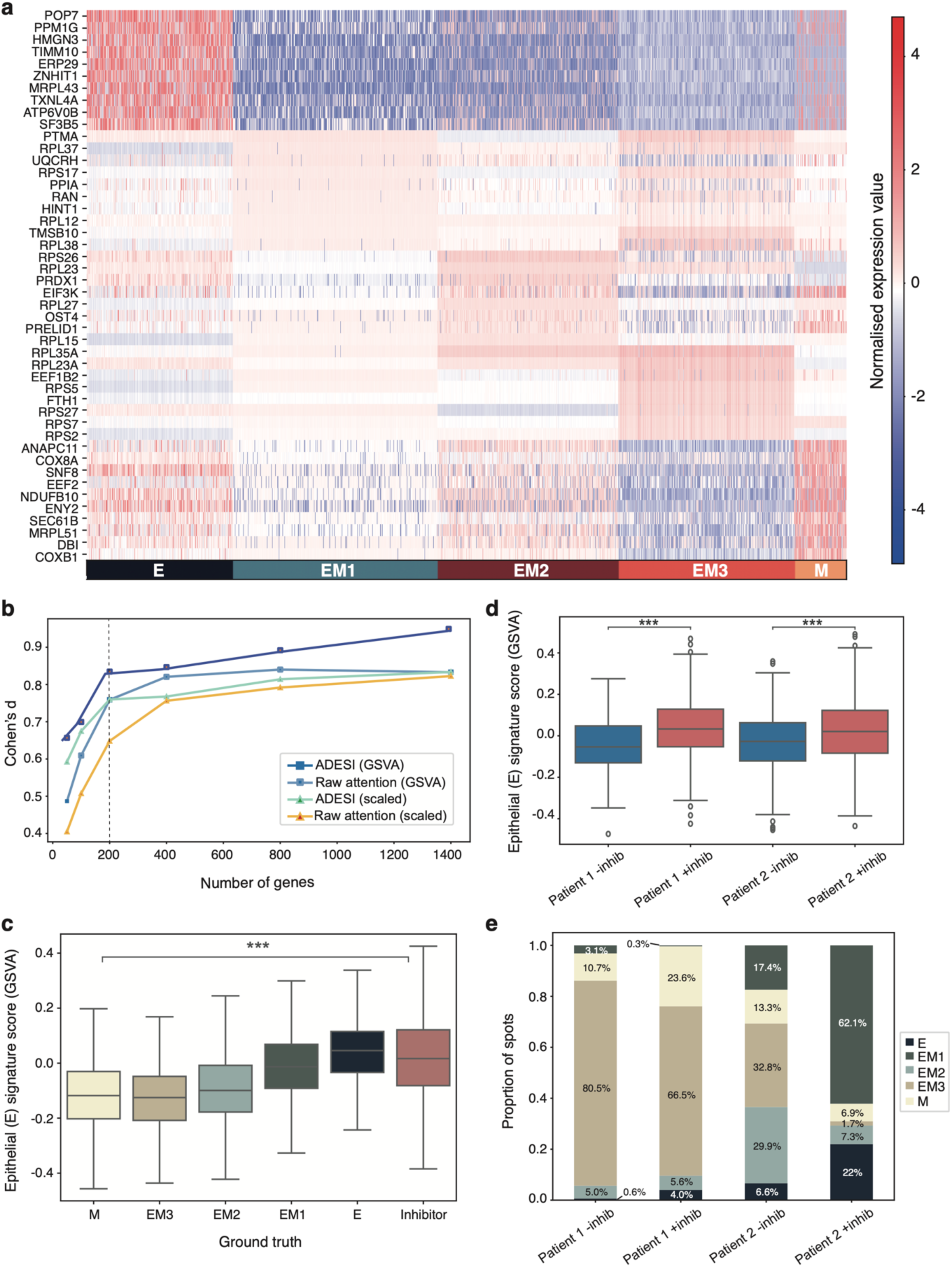
EMP gene signature inference and sensitivity analysis. **(a)** Expression heat map of key genes selected by the model using ADESI scoring across different categories, with a gradient from blue to red indicating increasing expression. The colour bar at the bottom distinguishes the EMP state for each profiled sample, offering an intuitive representation of which genes are important in predicting various EMP states. **(b)** Sensitivity analysis plot highlighting the effect of increasing the gene list size and alternating both methods to rank genes (raw attention vs. ADESI) and methods to score the gene list (scaled versus GSVA) on distinguishing the EMP states in the Cook and Vanderhyden^9^ data. The dashed line highlights the elbow point. **(d)** Validation of the epithelial gene signature in the EMT inhibitor (Netrin-1) dataset from Cassier et al^39^ (*** p<0.001). An increase in the epithelial signature is seen in both patients after the treatment with the inhibitor (+inhib).

Importantly, model-predicted scores exhibited different scales across datasets, reflecting complexities regarding EMT entry timing, duration and progression dynamics that lack established analytical frameworks.

### Gene programmes underlying TGFβ-induced EMT state transitions in cancer

To understand the gene regulation patterns underlying scFMs’ predictions, we developed the Attention-Driven Expression Significance Index (ADESI) to prioritise biologically meaningful changes that overcome the scFMs’ sensitivity to minor expression variation (see Methods). ADESI-identified genes showed distinct expression dynamics across EMT states in EMP-FM (**Figure 5a**), delineating these states more effectively than differentially expressed genes that exhibited extensive within-state heterogeneity (**Supplementary Figure 10**). Optimisation analysis (see Methods) revealed that a list of 200 genes provided sufficient information for reliable EMT state determination (**Figure 5b, Supplementary Table 6**). Validation using EMT inhibitor datasets from Cook and Vanderhyden^9^ and Cassier et al^39^ confirmed the gene list’s biological relevance, with epithelial (E) signatures increasing upon inhibitor treatment as expected (**Figure 5c-d**). The processes captured by the model included classical mesenchymal regulation markers such as *VIM* upregulation in EM3, but also other genes involved in processes linked with EMT such as oxidoreduction and ATP synthesis^40^, metabolic reprogramming, TGFβ signalling, motility and metastasis^41,42^ (**Supplementary Table 7**).

## DISCUSSION

In this work, we unveil the potential and limitations of applying scFMs as a tool for exploring cellular plasticity. Defining EMP as a prime example of a dynamic and plastic cellular process that involves multiple cell state switches, we systematically benchmarked scFMs against conventional machine learning and bioinformatics approaches through a comprehensive evaluation framework encompassing multiple datasets representing the same targeted underlying biological process. Our results demonstrate that, while scFMs show promise in capturing subtle gene regulatory patterns underlying EMP beyond up/downregulated genes inferred through traditional bioinformatics, significant performance disparities persist. Specifically, we identified that pre-trained models encode mixed signals from diverse biological processes, creating fundamental challenges when targeting complex phenomena like EMP that lack dominant transcriptional signatures - where subtle state transitions can be obscured by stronger tissue-type or cell-type signals.

Our results indicate that carefully designed adaptation strategies, such as the proposed dual-path classification/regression pipeline, can partially eliminate the gaps to achieve superior performance to simple fine-tuning pipelines and other SOTA methods. Our pipeline involved a naïve dual-path design with PEFT and MoE modules to systematically adapt pre-trained feature spaces while preserving foundational biological knowledge, enabling targeted optimisation for specific processes with reduced computational demands. Our proposed EMP-FMs can be considered as a proof-of-concept that scFMs, with careful design, could be adapted to model cell states and plasticity within limited settings. However, while our proposed approach shows some advantages, scFMs may underperform conventional methods in cross-dataset and domain shift scenarios. Substantial gaps remain for complex cell state classification tasks, and regression scores require recalibration. Thus, current scFMs require fundamental improvements in feature representation and architectural design to fully replace conventional approaches in future applications.

Compared to the cell type classification, predicting cell states is substantially more challenging due to their uncertainty. Single cell transcriptomic datasets offer time point snapshots during the process, with no guarantee that the states captured at those time points are stable. Conflicting evidence on EMP representing either continuous or discrete transitions further underscores this complexity^43–48^. This uncertainty demands models sensitive to subtle yet meaningful cellular perturbations - a requirement that distinguishes cell state classification from conventional cell type identification tasks. The proposed dual-path architecture addresses the challenge of capturing biological continuity from discrete, temporally sampled snapshots by jointly producing discrete state labels and continuous scores that reconstruct the underlying EMT continuum from potentially unstable cellular states. Continuous scores are preferable when experimental setups differ significantly from training conditions, but work to recalibrate these scores to new datasets will be required in the future.

Notably, the predictions from classical bioinformatics or machine learning methods aligned well with experimental measurements in several scenarios (e.g. spontaneous EMT) with much lower computational cost. Considering that, EMP is highly dependent on context^9,43^, both tissue and stimulus^9,43^, this emphasises the difficulty in developing a single unified framework for EMP and the value of traditional metrics for EMP alongside advanced foundation models, which potentially could be used in a complementary manner in the future. The fact that our EMP-FMs do not simply recapitulate classical EMT drivers but rather capture broader RNA metabolism, cellular energetics and motility features that accompany EMT, suggest their potential advantage over other methods in uncovering new regulators of this process, which will need to be validated experimentally. Future work should focus on expanding the capability of such models to capture a broader array of EMT programmes profiled under different stimuli and in more physiological contexts integrating the tumour microenvironment. However, such longitudinal datasets with ground truth EMT tracking are limited at the moment.

This study establishes a proof-of-concept that scFMs can be systematically adapted to characterize complex biological processes exhibiting continuous rather than discrete transitions, while also revealing persistent performance gaps relative to conventional approaches. By introducing a dual-path adaptation framework compatible with PEFT and MoE architectures, we outline a potential direction for extending pre-trained scFMs to other dynamic cellular processes. Our results indicate that fundamental improvements are still needed in scFM pre-training and architectural design, but scFMs hold potential in understanding biological complexity beyond conventional classification tasks.

## METHODS

### Training dataset

We sourced gene expression data from Cook and Vanderhyden^8^, made available at: https://drive.google.com/drive/folders/1SIEIf7UswTv_0S6TypYsaRzMcfkfsgji (GSE147405). The dataset encompasses 53,290 cells profiled through scRNA-seq from A549 (lung), DU145 (prostate), MCF7 (breast) and OVCA420 (ovarian) cancer cell lines, subjected to TGFβ (9,084 cells), TNF (22,159 cells) and EGF (22,047 cells) induced EMT, paired with EMT sampling times ranging from 0 day (epithelial state), 8 hours, 1 day, 3 days (hybrid states labelled EM1, EM2, EM3, respectively) to 7 days (closer to a fully mesenchymal state). These cell lines maintain an epithelial morphology and have been demonstrated to undergo EMT in previous studies. Morphological changes consistent with EMT in each cell line have been documented in Cook and Vanderhyden^9^. We employed raw counts as input to the model and followed the same preprocessing steps described in the protocol of the backbone model (scBERT) in all of our experiments. We also used the pre-computed pseudotime values originally generated by Cook and Vanderhyden^9^ as ground truth to train a regression model, enabling it to predict a cell’s position along this continuous spectrum.

In our training, we developed models using data from all tissue types, with specific models tailored for different stimulation conditions. Based on our training strategy, each stimulation type was associated with its own dedicated model, and we selected the best-performing models for subsequent validation experiments.

### Learning discrete cell states and pseudotemporal ordering by using a single cell foundation model with PEFT

We create a two-stage dual path framework for EMP task. In training stage 1, we trained a binary classifier for each EMP state, which served as an expert network in the following stages. This proxy task was designed to force the model to focus on a generalisable EMP process across different tissues. The expert network for each EMP state was adapted from scBERT^11^ with extra residual connections to enhance the raw gene expression signal and a Parameter-Efficient Fine-Tuning (PEFT)^21^ module. We named this expert network ‘scMultiNet’. Its purpose was to reshape the pre-trained feature manifold, redirecting the model’s attention towards more nuanced cellular state features relevant for EMP progression. EMP-FM involves multiple scMultiNets for different cell states, combined through a fusion network with Mixture of Expert (MoE)^22^ architecture. We simultaneously train a classification head for cell states and a regression head using the pseudotemporal ordering calculated by psupertime^9^ in the training set to capture the continuous nature of EMP (see Methods). This two-stage dual modelling approach enables our model to predict continuous scores that reflect EMT progression while preserving the integrity of the learned feature space.

Training stage II aims to train a fusion net to produce the final outcome. The gene expression profile data was filtered as required by scBERT and encoded into 200-dimensional vector spaces for each expert network (scMultiNet-E, scMultiNet-EM1, scMultiNet-EM2, scMultiNet-EM3, or scMultiNet-M). The selected pre-trained foundational model (scBERT or other potential single-cell foundation models in the future) is adept at analysing scRNA-seq data, aiming to generalise cell state predictions across various tissues and patient cohorts. As a pre-trained single-cell foundation model, the scBERT model underwent extensive pre-training on a corpus of approximately 100,000 scRNA-seq samples with gene2vec and a vocabulary of around 16,000 genes. The proposed method follows all the preprocessing steps in scBERT to make sure the proposed model can assign positional significance to genes.

We name our expert models ‘scMultiNet’ as we employ a simple multiplication layer consisting of fine-tuned foundation models. The proposed model considers the attention vector of the foundation model as a weight of raw gene expression profile data. We employed LoRA, a PEFT method, in our training stage I to adapt the embedding space foundation model’s architecture for interpreting single-cell gene expression profiles (**Figure 1**). PEFT enhanced the model’s adaptability to the targeted dataset while minimising the risk of overfitting. The design of scMultiNet ensured the model’s focus remained on genes relevant to the EMT process, filtering out those with low expression or tangential relevance. Genes exhibiting minimal expression were removed through normalisation and selective filtering to mitigate their potential to skew the model’s learning outcomes.

In all training stages, we utilised a one-dimensional convolution for dimensionality reduction. A three-layer neural network in training stage I is used to transform gene feature vectors into probabilistic cell type identities for binary classification proxy tasks. scMultiNet was trained with Adam optimizer (learning rate of 1e-4) and a cosine step scheduler.

For the target classification and regression tasks, we implemented a Mixture-of-Experts (MoE) architecture consisting of specialized expert mapper and a gating mechanism. Each expert mapper is structured as a two-layer perceptron with ReLU activation and dropout regularization. The input features are first processed through individual feature encoders, which project each feature group into a common representation space. A gating network, implemented as a multi-layer neural network, dynamically assigns input samples to the most relevant experts by computing importance weights. We employ a top-k gating mechanism with added noise during training to encourage diverse expert utilization. The expert outputs are then combined according to the gating weights, and the fused representations are processed through a final classifier/regression network comprising two fully connected layers with ReLU activation for the ultimate prediction tasks.

### Model validation

Validation of the EMP-FM model’s performance was carried out by randomly splitting the original data into training (80%) and testing (20%) sets, focusing on assessing the model’s accuracy. The model’s feature space was visualised using Principal Component Analysis (PCA) with Scanpy and Potential of Heat-diffusion for Affinity-based Transition Embedding (PHATE) with its original implementation from the Krishnaswamy lab^23^ to compare the changes before and after training, demonstrating that the training process effectively captured the key information related to EMT.

During training, we randomised the order of all sampling time points and discretised them into one-hot encodings. By not introducing temporal information during training, if the learned embedding space exhibited clear time-series associations, it would indicate that the model had learned the correct information (results shown in **Figure 2 and Supplementary Figure 2**).

To further validate the robustness and broad applicability of our approach, we tested our model on independent single-cell datasets where EMT is tracked longitudinally, with different experimental designs. These datasets are available for download from GSE194019, GSE114687, GSE155628 and GSE149299. The model’s performance was analysed by comparing the categories provided by the datasets with the continuous scores inferred by the model.

In our analysis of the GSE155628 and GSE149299 datasets, we mapped mouse genes to human homologues using Ensembl BioMart. After data cleaning and integration with the gene annotation file, approximately 80% of mouse genes were successfully mapped to identifiable human homologues. For genes with multiple human orthologues, we used an averaging approach to consolidate expression values.

### Comparison with traditional machine learning and bioinformatics methods

The evaluation process involved an initial dimensionality reduction via PCA across the first 10 PCs while retaining 95% of variance, followed by training with machine learning algorithms (KNN, random forest and AdaBoost from SciPy). We also conducted comprehensive comparisons against two categories of baseline methods. The first category includes state-of-the-art single cell foundation models: scGPT, scFoundation, and Universal Cell Embedding (UCE), which were fine-tuned on our training data. Conventional bioinformatics analysis methods (scType^29^, AUCell^28^ and scPred^27^) were also tested to demonstrate a full comparison with all potential state-of-the-art methods. For fair comparison, we implemented two distinct evaluation frameworks. On the Cook dataset, all methods were evaluated using identical train-test splits, with performance measured by the average Area Under the Receiver Operating Characteristic curve (AUROC) for EMT state classification. For additional independent validation datasets where discrete states and temporal sampling designs differed from the training data, we adopted a continuous scoring approach, evaluating performance through Pearson Correlation Coefficient (PCC) between predicted EMT scores and experimental sampling timepoints. All methods share the same training set.

To identify differentially expressed genes (DEGs), we applied the FindAllMarkers function in Seurat. Marker genes were identified by comparing cell populations defined by the ground truth annotation on the Cook and Vanderhyden^9^ full dataset. We focused on genes that were upregulated in each group by setting only.pos = TRUE. To ensure robust detection, we applied a minimum expression threshold (min.pct = 0.5), requiring that genes be expressed in at least 50% of cells in each group. Additionally, we set a low log-fold change threshold (logfc.threshold = 0.1) to capture subtle expression differences relevant to state transitions. Then, we applied AUCell scoring using DEGs for each category as one mega list, which performed better than using individual gene lists for every time point. Results were visualised as estimation plots^49^ to demonstrate median differences from baseline category.

### Attention-Driven Expression Significance Index (ADESI) scoring

We introduce the Attention-Driven Expression Significance Index (ADESI), a novel method designed to enhance the interpretability of complex biological datasets. Traditional single-cell foundation models use attention visualisation to demonstrate model interpretability and versatility. However, this may fail to capture changes between states that would be considered impactful enough to be biologically meaningful. ADESI addresses this issue by leveraging the EMP-FM architecture and Z-scores to provide biologically relevant gene weightings. Initially, mirroring traditional methods, we extracted attention scores from each expert model relevant to the designated category. These scores were then multiplied by the original expression levels of the respective genes to derive the EMF-FM output. The average score for each gene was calculated in the final classified category and the top K genes were prioritised. This design is used to aid researchers in understanding the genes that the model’s attention mechanism focuses on, which could be potentially employed as markers of these states.

### Spatial transcriptomics analysis

Endometrial carcinoma Visium slides were obtained from Cassier et al^39^. Filtered matrices were loaded, merged per patient and spots with fewer than 1,000 detected genes were removed. Pre-processing and normalisation were conducted using the Scanpy package. To more precisely identify tumour cells we employed the copy number inference tool STARCH^50^ and only kept spots that showed evidence of copy number changes, which are likely to be tumour specific. SpottedPy^51^ was used to identify EMT niches and assess the spatial relationships.

### Gene signature analysis

We used Gene Set Variation Analysis (GSVA) calculated using the GSEApy package^52^ to score the tumour samples with the EMP-FM gene signatures. We categorised the high attention genes based on their upregulation or downregulation in the initial dataset for each time point. We calculated a composite score by subtracting the GSVA score of downregulated genes from that of the upregulated genes. The scaled score for comparison was calculated by subtracting the Z-score of downregulated genes from the Z-score of the upregulated genes. Cohen’s d score was used for assessing how well the gene sets distinguish each cluster. Cohen’s d was computed by assessing the difference in means between the groups, standardised by their pooled variability. G:profiler^53^ was used for gene set enrichment analysis.

### Model building, statistical analysis and data visualisation

The EMP-FM model was built with PyTorch for transformer blocks, and pytorch-lightning for efficient training. The analyses of the results were conducted in Python and R. Groups were compared using a two-sided Student’s t-test, Wilcoxon rank-sum test or ANOVA as required. Correlations were calculated using Spearman’s rank order correlation. Graphs were generated using the Seaborn and Matplotlib Python packages.

## DATA AVAILABILITY

The results published here are based upon publicly available data made available at the Gene Expression Omnibus (GEO) by Cook and Vanderhyden^9^ (GSE147405), Paul et al^35^ (GSE194019) and McFaline-Figueroa et al^36^ (GSE114687). The endometrial cancer spatial transcriptomic slides were downloaded from Cassier et al^39^ (GSE225691). Single cell RNA-seq data of the adult mouse mammary epithelium from control and Zeb1 MSKO mice were downloaded from Han et al^38^ (GSE155628). Single-cell RNA sequencing data from C3(1)-Tag cancer cells tracked during invasion and colony formation were obtained from Grasset et al^37^ (GSE149299).

## CODE AVAILABILITY

The EMP-FM package can be found at the following GitHub repository: https://github.com/secrierlab/EMP-FM. The implementation of the basic module of EMP-FM, namely scMultiNet, can be accessed within this repository at: EMP-FM/scLLM/Models/scMultiNet. All the code developed for the purpose of this analysis is deposited at: EMP-FM/Experiment.

## AUTHOR CONTRIBUTIONS

MS and SP designed the experiment. MS supervised the analyses. SP developed and tested the EMP-FM model and the ADESI score. EW prepared the training and validation datasets, performed the gene sensitivity analysis, pathway enrichment and spatial transcriptomics analyses. CC evaluated EMT in validation datasets using classic EMT markers and bioinformatics methods. All authors wrote and approved the manuscript.

## ACKNOWLEDGEMENTS

MS, SP and CC were supported by a UKRI Future Leaders Fellowship (MR/T042184/1, MR/Y034031/1). EW was supported by a studentship award from the Health Data Research UK-The Alan Turing Institute Wellcome PhD Programme in Health Data Science (218529/Z/19/Z). Work in MS’s lab was supported by a BBSRC equipment grant (BB/R01356X/1) and a Wellcome Institutional Strategic Support Fund (204841/Z/16/Z).

The illustrations in Supplementary Figure 1 were generated using the icon libraries available at https://bioicons.com/.

## ETHICS DECLARATIONS

This study employs only publicly available data. All data comply with ethical regulations, with approval and informed consent for collection and sharing already obtained by the relevant consortia.

## SUPPLEMENTARY TABLE CAPTIONS

**Supplementary Table 1.** Performance comparison of different state-of-the-art models for EMT prediction tasks on the datasets from Cook et al. The table presents AUROC scores for various models across three stimulation conditions (TGFβ, EGF, and TNF) and their averaged performance. Models are categorized into three groups: our proposed EMP-FM models with different parameter sizes (512 and 128), state-of-the-art foundation models (SOTA-FM including scGPT, Geneformer, and UCE), and conventional machine learning approaches (random forest, KNN, and AdaBoost). Higher scores indicate better predictive performance.

**Supplementary Table 2.** Performance of the three EMP-FM models (columns) in predicting EMT states across the different stimulation datasets (listed as rows).

**Supplementary Table 3.** Performance of the three EMP-FM models in predicting each EMT state (timepoint) in each cancer type.

**Supplementary Table 4.** Kruskal-Wallis test statistics showcasing group differences for individual figure panels where EMP-FM regression scores as well as AUCell-generated scores based on DEGs and classical EMT markers are compared in validation datasets.

**Supplementary Table 5.** Performance of EMP-FM, scGPT and random forest regression models in the external validation datasets. The top performing model in each dataset is highlighted in red. PCC - Pearson correlation coefficient.

**Supplementary Table 6.** The top 200 up-/down-regulated genes in each EMT state, as uncovered by the ADESI metric and robustness analysis.

**Supplementary Table 7.** Pathway enrichment analysis results for the EMT gene signatures. **(a)** The 0d (E) gene signature. **(b)** The 8h (EM1) gene signature. **(c)** The 1d (EM2) gene signature. **(d)** The 3d (EM3) gene signature. **(e)** The 7d (M) gene signature.

