## Supplementary Material for "From cell types to cell states: adapting foundation models for cellular plasticity"

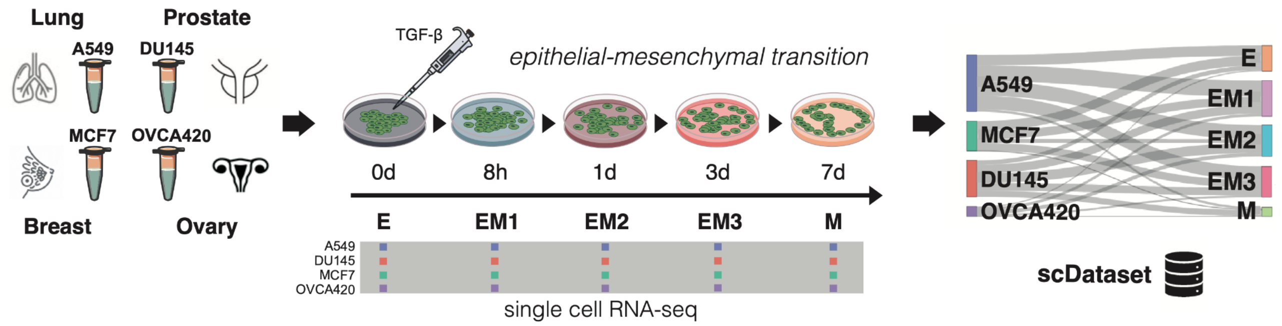


**Supplementary Figure 1: Experimental setup for the EMT training data**. scRNA-seq profiles of lung (A549), prostate (DU145), breast (MCF7) and ovarian (OVCA420) cell lines before EMT induction with TGFβ, TNF and EGF (0 days - 0d) and 8 hours (8h), 1 day (1d), 3 days (3d) and 7 days (7d) after stimulus, respectively, are employed as a ground truth dataset for EMT transformation.

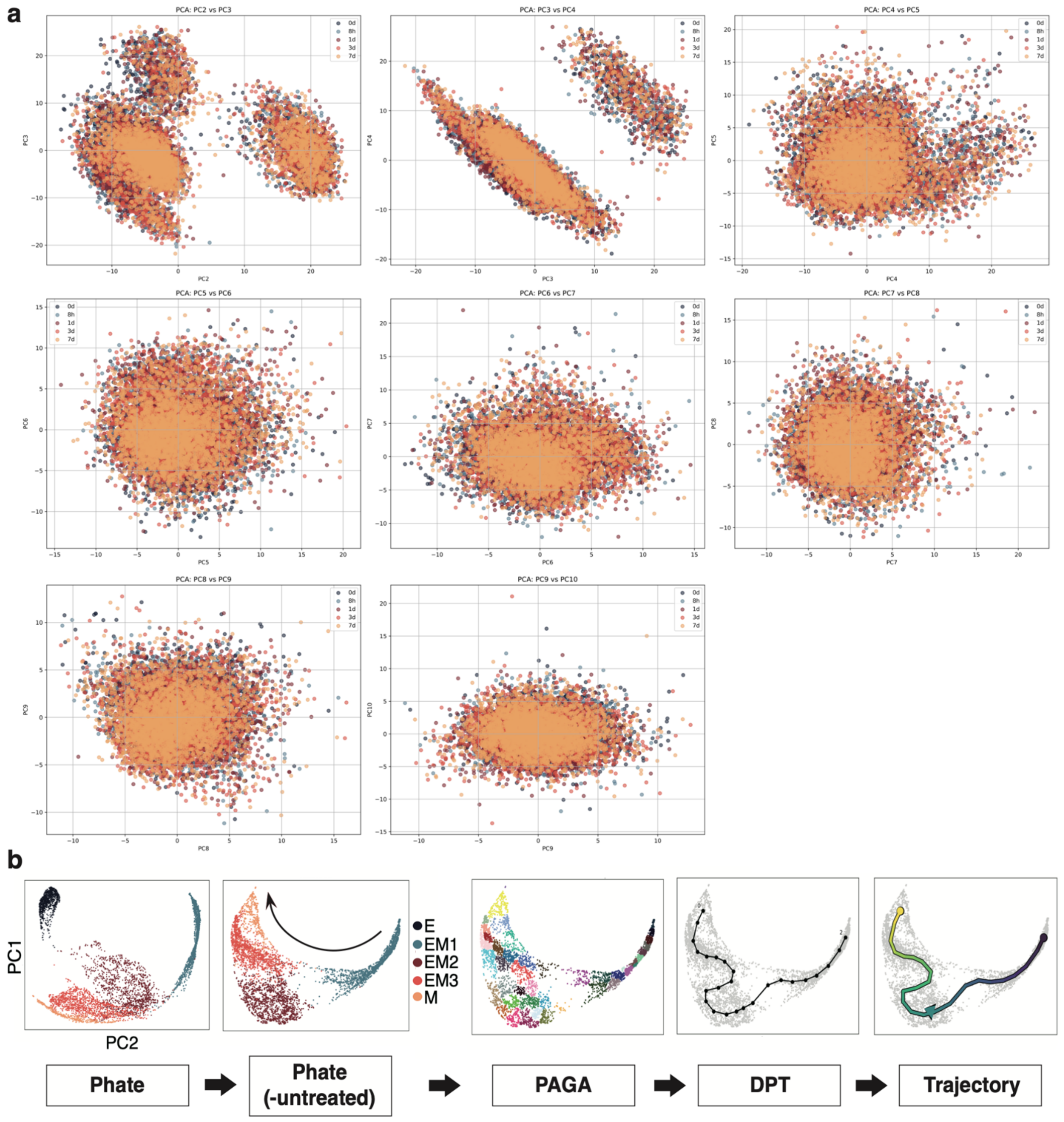


**Supplementary Figure 2: Projections of the gene expression profiles and trajectory analysis. (a)** PCA projections denoting principal components 2-10 of the raw gene expression profiles are plotted in a pairwise fashion. Every dot denotes a cell, coloured according to the time point when it was sequenced, corresponding to distinct EMT states (0d - E, 8h - EM1, 1d - EM2, 3d - EM3, 7d - M). No PCs capture variation that is directly linked with the EMT state. **(b)** PHATE visualisation emphasises the EMT trajectories captured by the embedding space of the final EMP-FM model. From left to right: visualisation of all sampled data points using PHATE, only samples after TGFβ stimulation (removing day 0, arrow indicates the time course progression), PAGA, DPT and trajectory reconstruction. The PAGA step delineates the data into distinct clusters, which are then sequentially ordered using DPT to infer developmental trajectories, culminating in a trajectory plot that captures the dynamic journey of cell fate decisions.

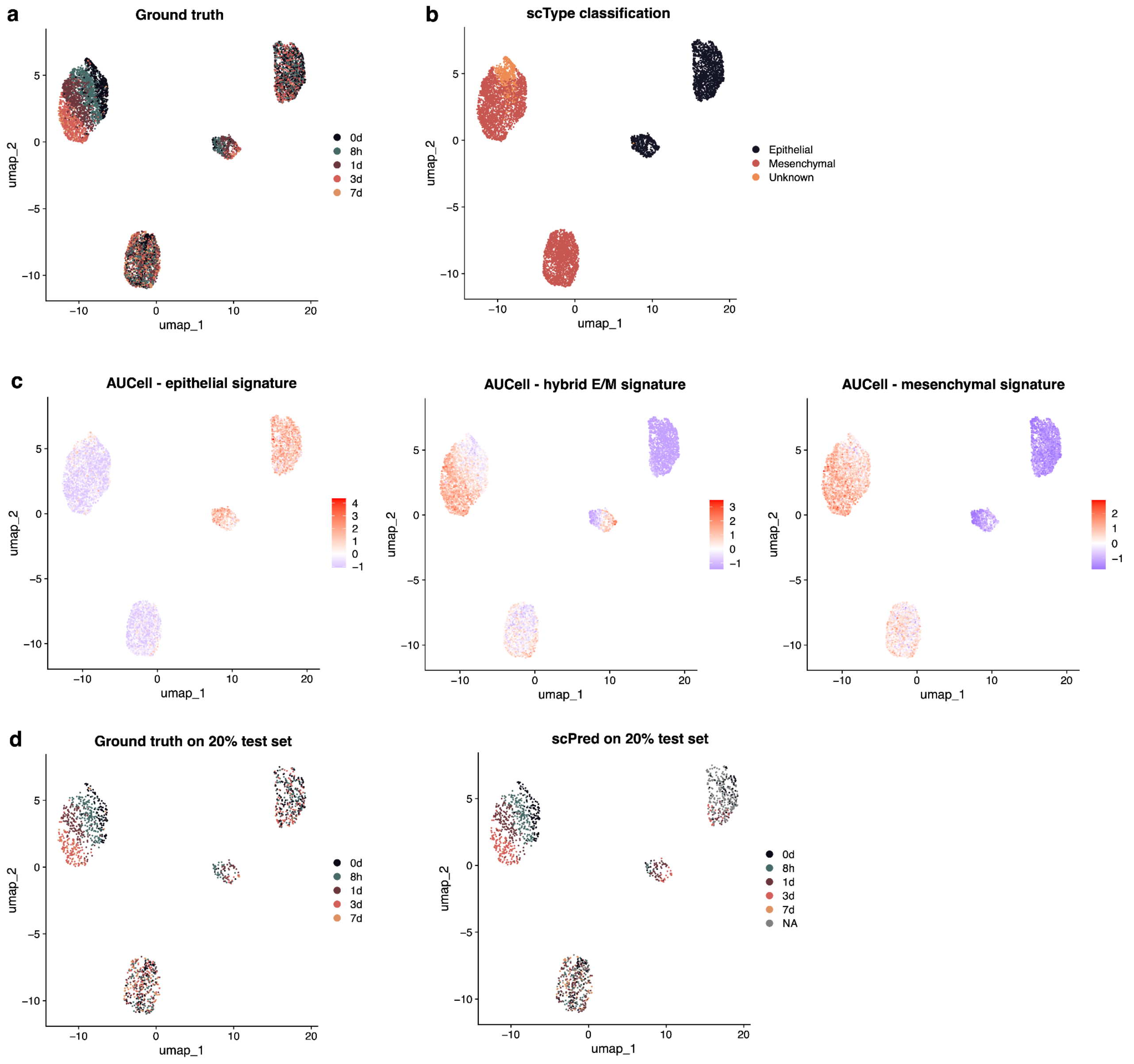


**Supplementary Figure 3: Evaluation of conventional bioinformatics methods for predicting EMP. (a)** Ground truth labels of the Cook and Vanderhyden full dataset. **(b)** scType annotations based on EMT gene lists on the Cook and Vanderhyden dataset. **(c)** AUCell scores derived from EMT gene lists in the Cook and Vanderhyden dataset, illustrating epithelial (left), hybrid (middle) and mesenchymal (right) states. **(d-e)** Ground truth labels (d) and scPred predictions (e) on a 20% test dataset, with training performed on 80% of the Cook and Vanderhyden dataset.

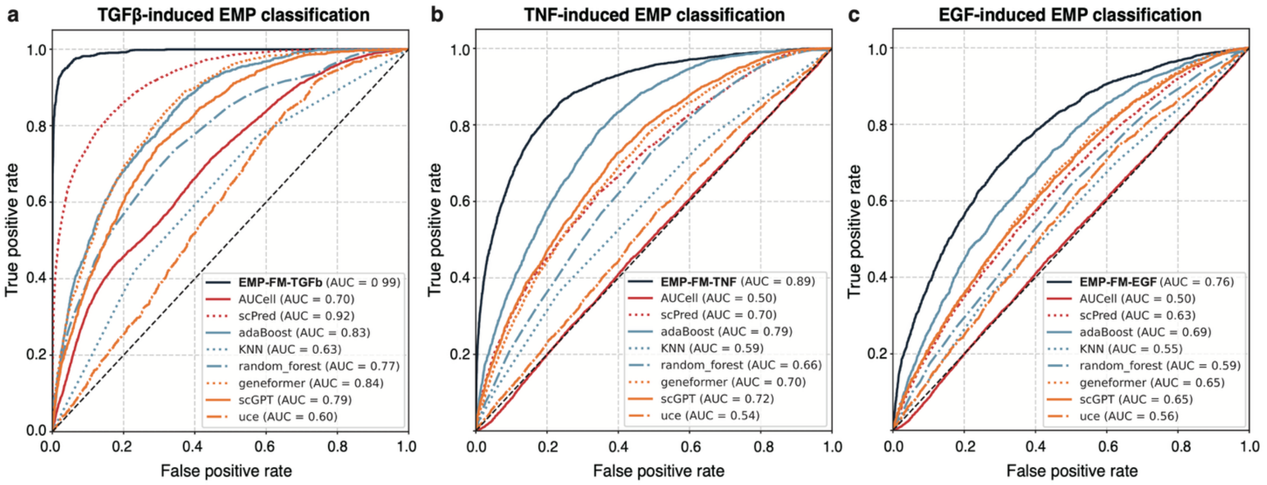


**Supplementary Figure 4: Performance comparisons across different EMP-FM models and SOTA methods**. The performance of stimulus-specific models: **(a)** EMP-FM-TGFβ, **(b)** EMP-FM-TNF and **(c)** EMP-FM-EGF is compared against that of various foundation models (Geneformer, scGPT, UCE), conventional ML approaches (AdaBoost, KNN, random forest) and bioinformatics methods (scPred, AUCell).

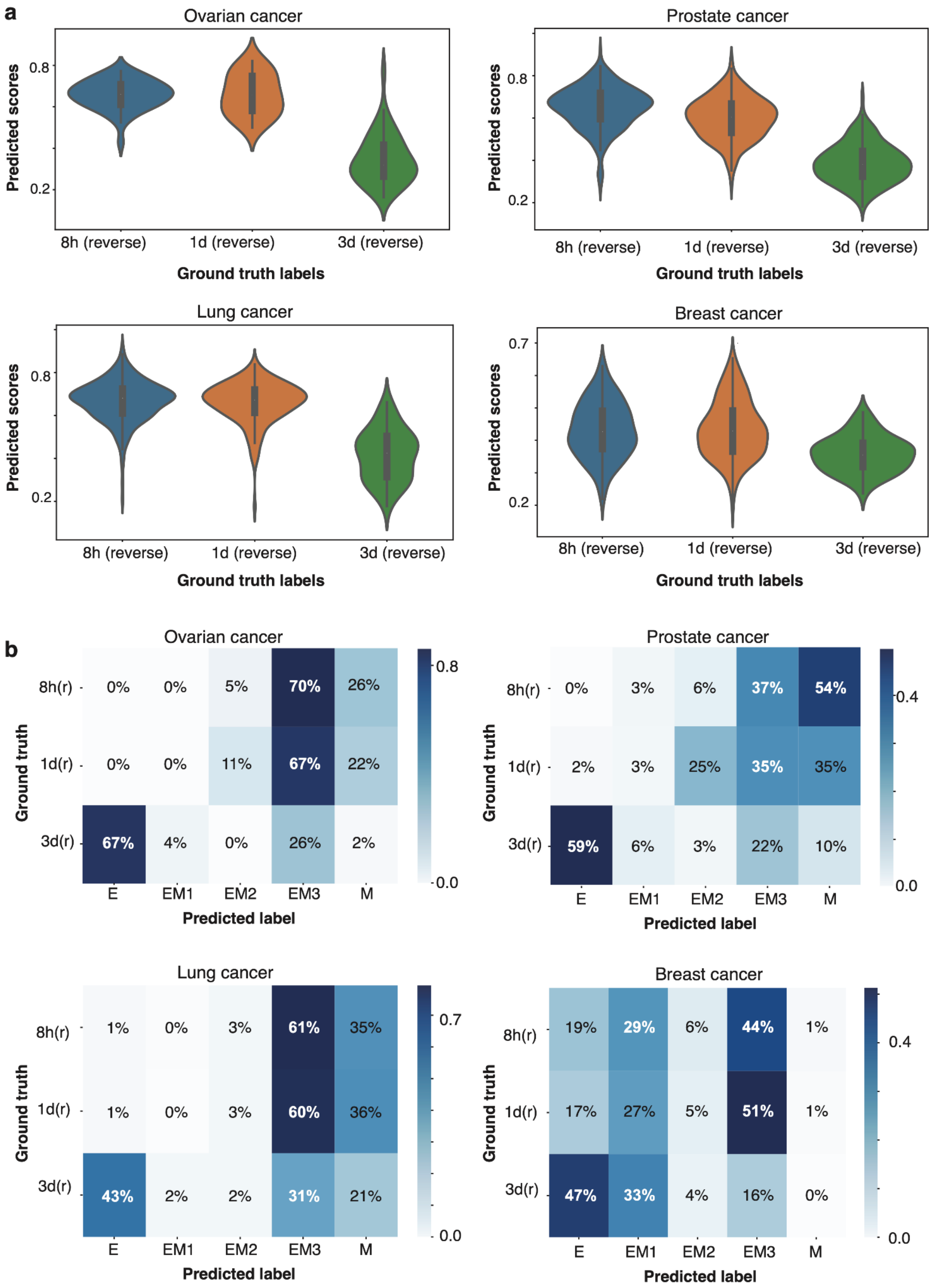


**Supplementary Figure 5: EMP-FM predictions in the TGFβ stimulus removal data in every cancer cell line. (a)** EMP-FM-TGFβ predicted regression scores for ovarian, prostate, lung and breast cancer cell lines upon reversal of the EMT process, measured at 8 hours, 1 day and 3 days after stimulus removal. **(b)** The confusion matrices for different cancer types, contrasting ground truth categories and categories predicted by the EMP-FM-TGFβ model.

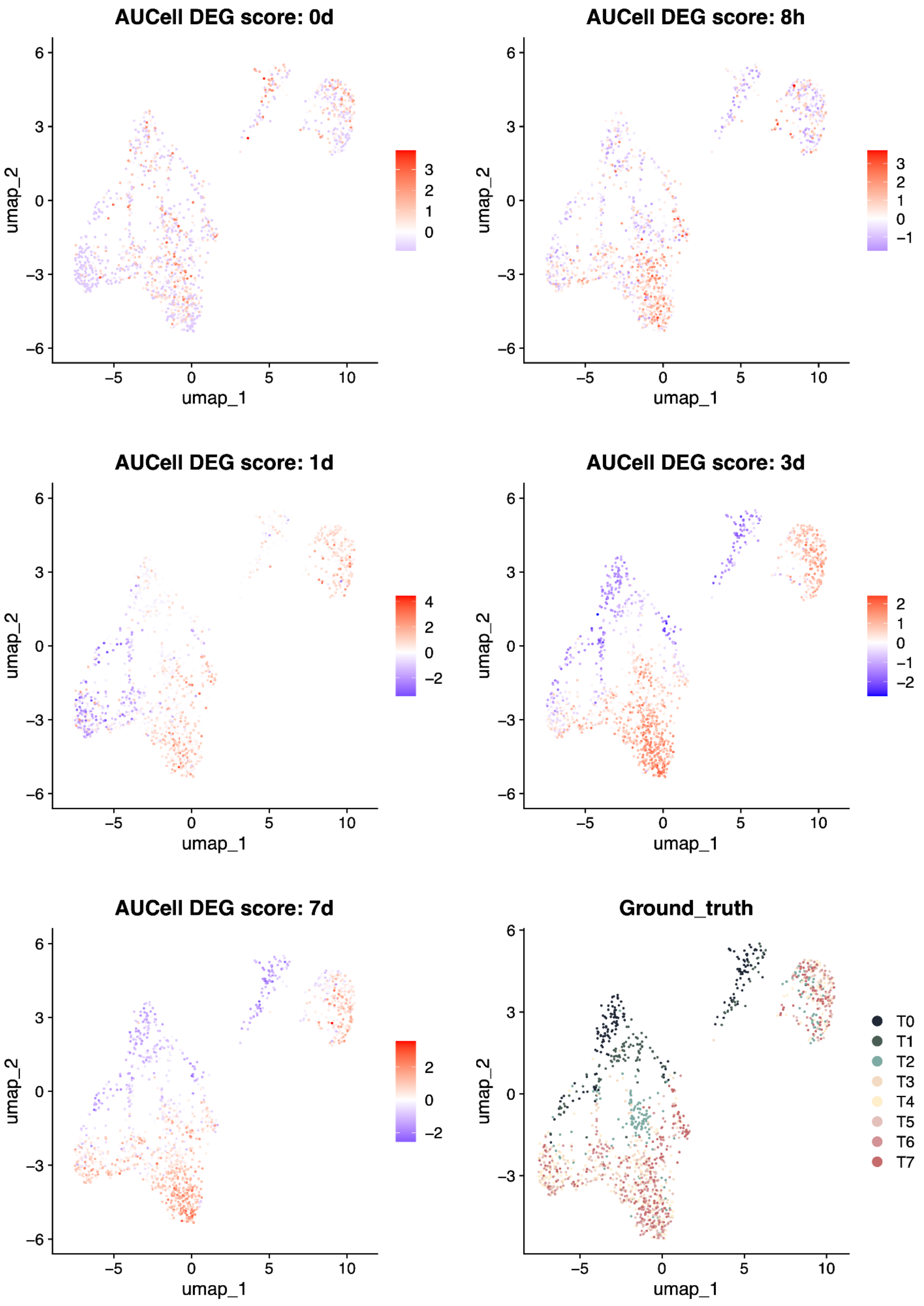


**Supplementary Figure 6: Applying differentially expressed gene lists to identify EMT states in the Paul et al dataset.** AUCell calculated scores per cell based on differentially expressed genes at distinct time points in the Cook and Vanderhyden dataset are shown applied to the Paul et al time points. The ground truth time point labels are also shown (bottom right).

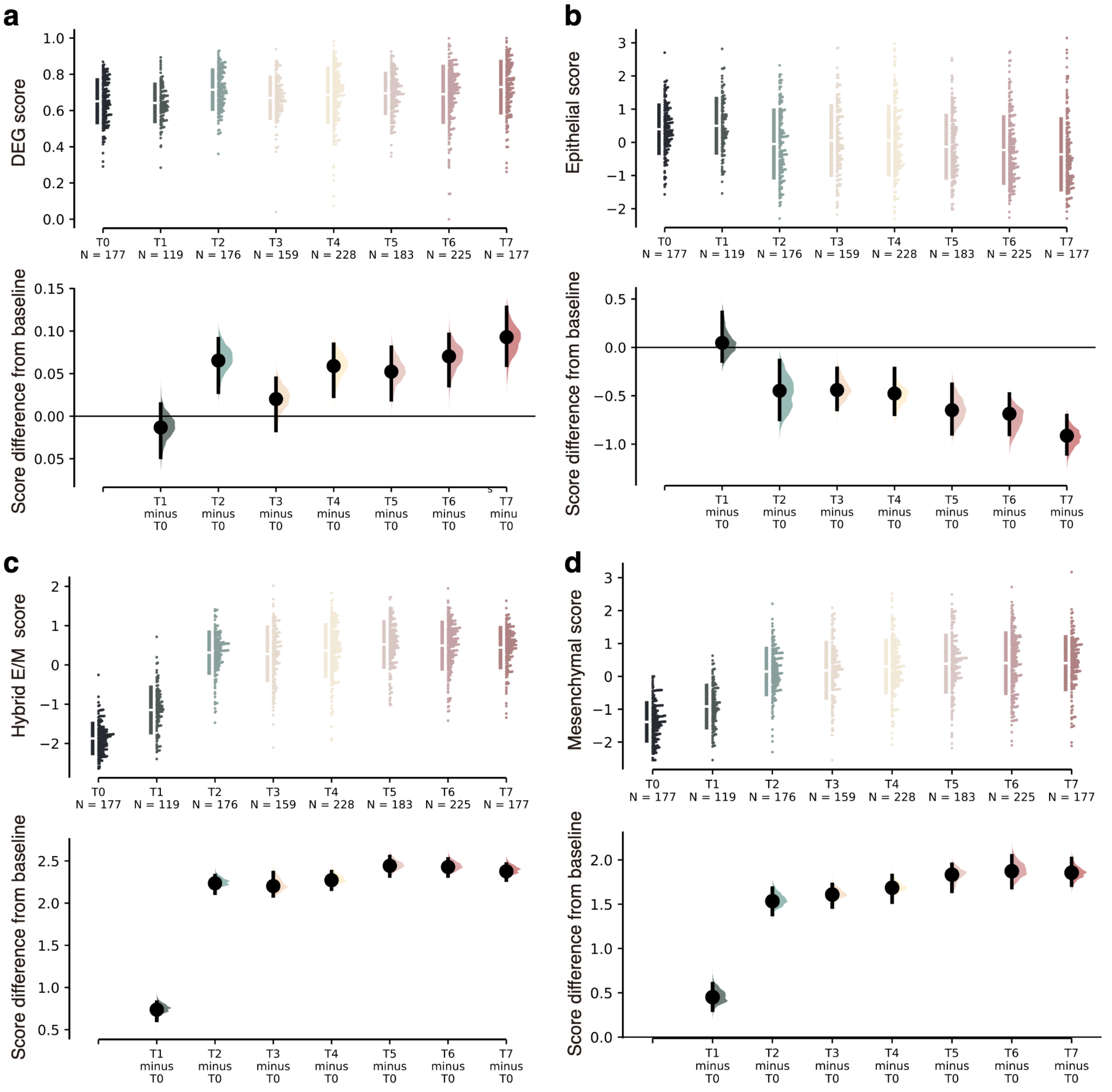


**Supplementary Figure 7: Validation of conventional EMT scoring strategies in the longitudinal dataset from Paul et al.** Predicted scores per time point (top) and difference in scores compared to the baseline, i.e. T0 (bottom), are shown for the following gene signatures: **(a)** differentially expressed genes (DEGs) derived at each time point in the Cook and Vanderhyden dataset; **(b)** classic epithelial markers; **(c)** classic hybrid EMT markers; and **(d)** classic mesenchymal markers.

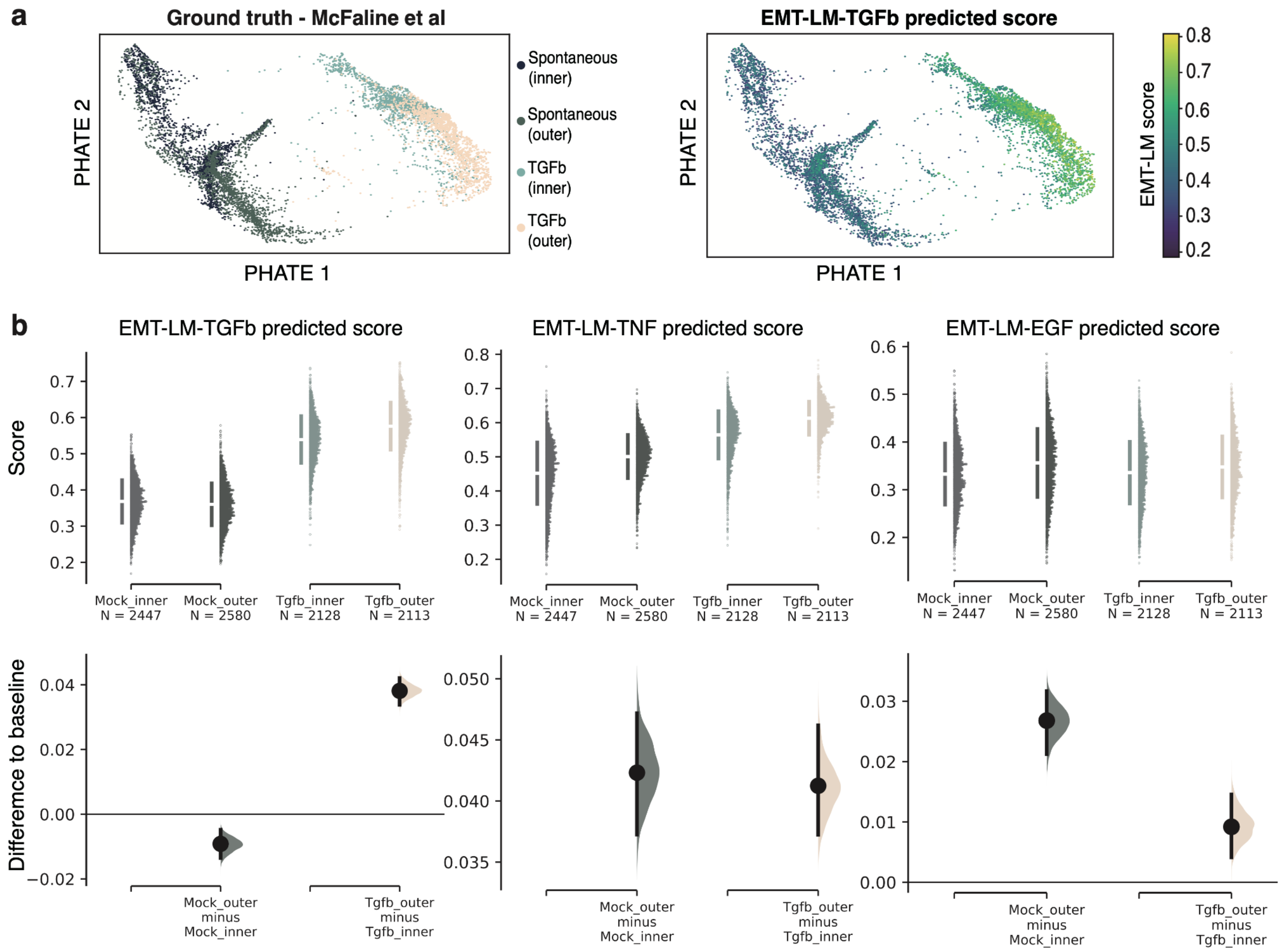


**Supplementary Figure 8: Validation of the EMP-FM-TGFβ model in the McFaline-Figueroa et al dataset. (a)** Left: PHATE visualisation of individual cell projections coloured by EMT stimulus (spontaneous or TGFβ-induced) and ground truth sampling time point (inner – before EMT transformation; outer – after EMT transformation). Right: Regression scores predicted by EMP-FM-TGFβ , with higher values indicating a more mesenchymal state. **(b)** Distribution of EMP-FM-TGFβ scores predicted for individual cells in the two stimulus conditions and before/after the EMT induction (inner-outer), alongside classic epithelial, partial EMT (pEMT) and mesenchymal scores from the literature. Violin plot colours are the same as in panel a.

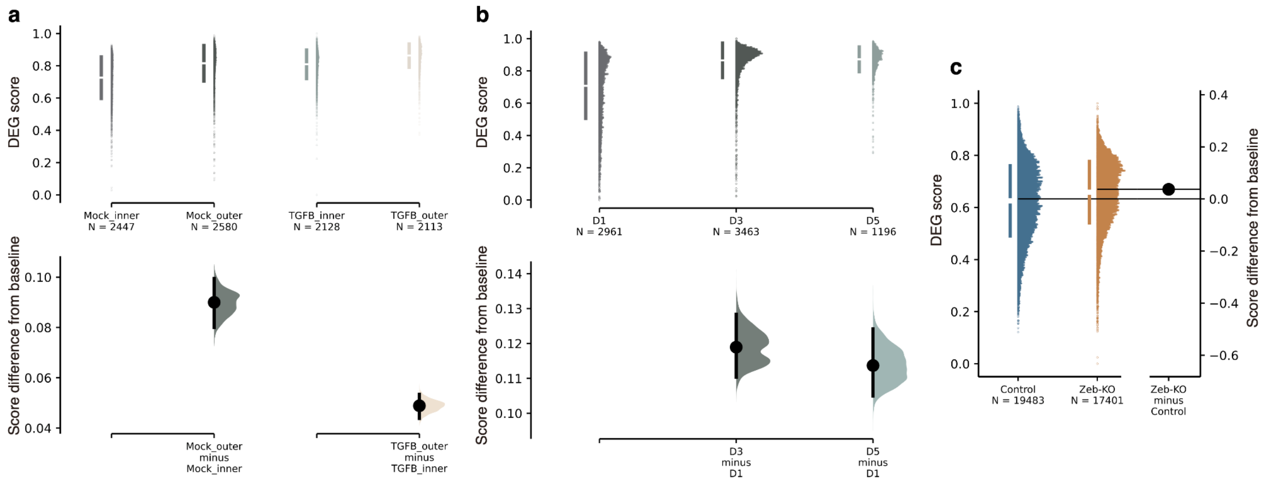


**Supplementary Figure 9: Differential gene expression signature performance in external datasets.** (a) DGE scoring in MCF10A cell lines undergoing EMT spontaneously (mock) or upon TGFβ stimulation (TGFB); inner – before EMT transformation; outer – after EMT transformation. The score distributions are shown in the top plot and the difference compared to epithelial state in the bottom. (b) DGE score distribution in C3(1)-Tag mouse breast cancer cells tracked during hybrid E/M transition (Grasset et al) at day 0 (12h), day 3 and day 5 (top) and differences in scores compared to the day 0 baseline (bottom). (c) DGE score distribution in mouse basal mammary epithelial cells (Han et al) tracked before and after Zeb1 knockout (left) and score difference compared to control (right). Changes compared to baseline are significant (Kruskal-Wallis test p<<0.001, see statistics in Supplementary Table 4).

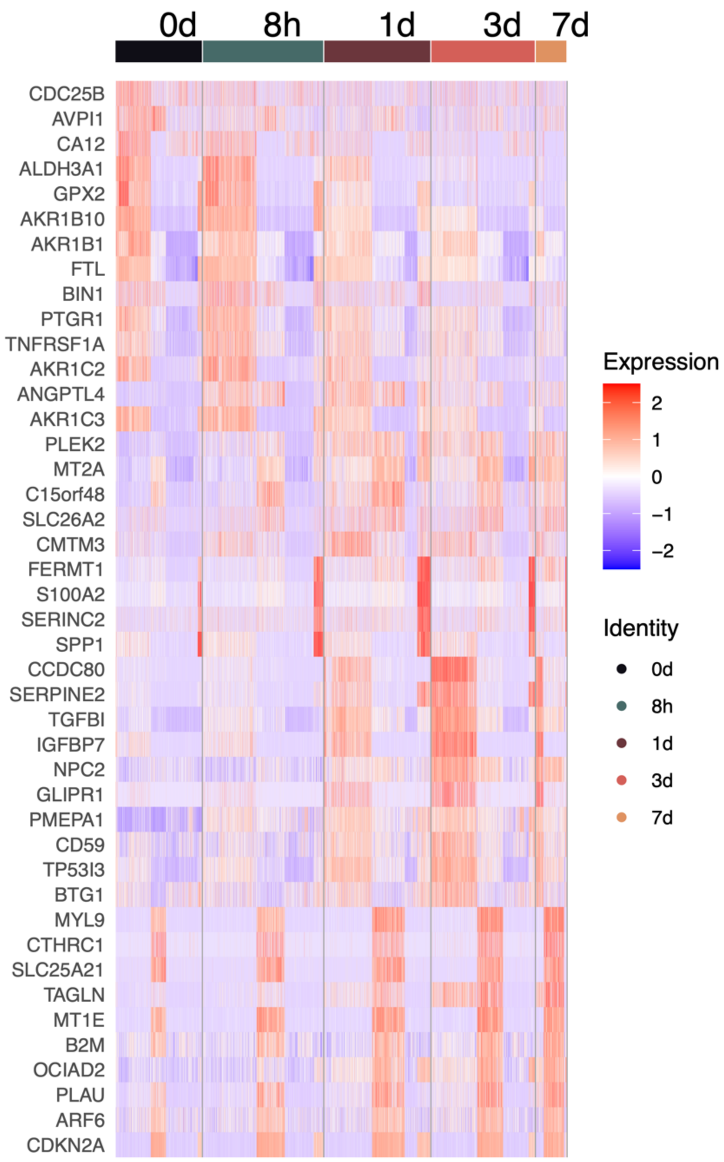


**Supplementary Figure 10: EMT state-specific expression patterns for differentially expressed genes at distinct time points.** The list of differentially expressed genes (rows) has been derived in the Cook and Vanderhyden dataset. Columns correspond to individual cells sequenced at distinct time points (in distinct EMT states).
